# Complexity of EEG Reflects Socioeconomic Context and Geofootprint

**DOI:** 10.1101/125872

**Authors:** Dhanya Parameshwaran, Tara C. Thiagarajan

## Abstract

The fine scale structure and resulting activity of the brain are largely shaped by experience, suggesting that the faster rate and complexity of experience offered by modern civilization may have significant impact on human brain dynamics. Here we defined a new measure of complexity of the EEG signal and compared it across populations spanning incomes from <$1/day to ∼$410/day with a wide range of access to features of modern life such as urban environments, higher education, electricity, motorized transport and telecommunication. Complexity across our sample spanned a 2.75-fold range, separating into two distinct distributions of pre-modern and modern experience. Furthermore, complexity scaled systematically with various technologies and experience factors, of which travel or geofootprint had the strongest relationship. Complexity also had a steep non-linear relationship with income that leveled out at an income of ∼$30/ day. Finally, it was strongly correlated to performance on a pattern completion task indicating its relevance as a cognitive measure. In light of growing income inequality and divergence in access to the tools of modern living across the globe, our findings have significant implications for social policy.

## INTRODUCTION

Until 1800 the majority of the world lived in small rural settlements, education and literacy were limited, and mobility was constrained by horse speed. The world has changed profoundly over the last 200 years with increasing agglomeration into large urban settlements alongside greater mobility and communication enabled by technology, and an explosion of collective knowledge. These elements of modern civilization greatly alter both the rate and the possible complexity of experience but have not perpetuated uniformly across the world. With incomes now diverging well over a million fold [1; 2] people increasingly diverge in their access and use of these tools, and therefore the rate and complexity of their experience of the world. What is the impact of this dramatic divergence on the human brain?

Unlike any other organ, the brain is profoundly experience dependent in its function. The precise circuitry and dynamics of the brain are sculpted by experience [3; 4; 5; 6; 7; 8; 9; 10] and changes in the level of activity result in not just quantitative, but also qualitative changes in the mechanism of neural function [11; 12]. Furthermore, studies done in rodents show that placing the animal in a more complex or enriched environment has far reaching impact on gene expression in the brain [13], the degree and nature of synaptic plasticity [14; 15; 16; 17; 18], dendritic branching[19; 20], brain surface area and a host of other functional and structural aspects [21; 22]. Enriched environment also protects against cognitive deficits in aging [23] and those resulting from other diseases such as epilepsy [24], Alzheimer’s[25] and Down Syndrome [26; 27; 28]. Furthermore, studies in the United States demonstrate that income, particularly childhood poverty, has a dramatic negative effect on cortical volume [29] and surface area [30], as well as gamma power in the EEG [31]. Conversely years of education have been shown to correlate positively with these structural aspects of the cortex [32; 33].

These findings strongly suggest that the advances of modernity may have dramatically altered the dynamical functioning of the human brain. We therefore address here how differences along core dimensions of income and access to the dominant tools and structures of modern civilization have impacted the complexity of dynamics of the human brain.

We chose to carry out this study in India, where modern cities exist in relatively close proximity to remote rural settlements with little access to modern tools. In order to restrict the impact of racial genetics and language, we confined our study to the State of Tamil Nadu, which is relatively homogeneous along these dimensions. For this study we collected resting EEG activity from participants for 3 minutes with informed consent in a quiet environment with their eyes closed, along with a host of demographic and socioeconomic information including income, education, occupation, travel patterns, mobile phone usage, electricity and fuel consumption and Internet usage. Our sample comprised 402 adults between the ages of 21 and 65 from 48 locations including remote settlements of just 300 people with no access to electricity or motorized transport, to cities of several million people with all modern amenities (see Methods, Sup. Figs. 1A,B.) Annual incomes of participants ranged from $300 to approximately $150,000 dollars, (conversion at 2014 exchange rates of Rs. 60 INR per USD) translating to daily incomes of $0.85 to ∼$410 (Sup. Fig.1C). The ∼483 fold range of incomes of our sample has a distribution that is roughly similar in structure to world income distribution, relevant to upwards of 90% of the global population. Across this range of incomes, formal education levels spanned anywhere from no schooling to college graduates, with college educated representing ∼10% of the sample, again in line with global metrics for education [34] (Sup. Fig.1D).

Our results indicate a distinct shift in complexity that is systematically related to signatures of modernity with significant implications for how we understand ourselves collectively and how we approach social and economic development around the world.

## MATERIALS AND METHODS

### Field Recruitment, Survey and Recording

Participants were recruited from 48 locations across the state of Tamil Nadu in India (Sup. Fig. 1A). Locations were selected to span various criteria of population, education levels, industrialization and infrastructural features such as distance to road, electrification, industrial worker ratio and other factors derived from the census, economic census and other geospatial data sources.

Participants were selected to span an income range from $0.85 to ∼$410/day (family income) and within each income band ($0-$10/day, $10-$30/day and >$30/day) were spread roughly equally by age and gender.

Willing participants who met our demographic sampling criteria (gender, age, income, no known history of physical or mental illness) were first surveyed to capture information about their income, education and technology use. Participants were then instructed to wash and dry their hair on the day of the EEG recording without the application of hair products, particularly hair oil, which is customary in the region. Low-income participants were provided with a sachet of shampoo.

Recording locations were often outside in rural locations and care was taken to select locations at distance from noise producing equipment such as mobile towers and electric motors and pumps. Low-income participants were sometimes paid Rs. 150 ($2.50) to compensate for loss of wages when experiments were conducted during working hours. Prior to or following the recording (but on the same day), participants were asked to participate in a simple test of memory recall. A subset (28 people in two locations spanning a range of complexity) was asked to complete a short pattern completion test.

All participants were fully informed about the intent and methodology of the experiment and signed a consent form. All recruitment, consent and data collection were carried out in accordance with protocols approved by Health Media IRB (USA, OHRP IRB #00001211) and Sigma-IRB (India) in accordance with Title 45, code of federal regulations, sub-part A of NIH and Indian Independent Ethics Committee requirements.

### Survey of Individual Experience

To understand the broad contours of how individuals across different socioeconomic contexts experience the world, our survey probed individual and family income levels, years of education, geo-footprint as measured by how far they had traveled in the world, geographic distance of their social network of friends and family, and finally use and spending on various modern technologies from fuel to electricity, mobile phones and the internet (Sup. Table 1).

### Cognitive Tests

#### NAME RECALL

Participants who had writing literacy (n=346) were asked to write down on a piece of paper as many names of people or places that they could think of in a one minute period. We note that this exercise was done prior to or immediately after the recording and not during where they remained still with their eyes closed.

#### PATTERN COMPLETION

The pattern completion test was an abridged adaptation of the Raven’s test and involved 5 questions of increasing difficulty. Questions were of a standard format showing a pattern and a multiple-choice selection of five options of which one completed the pattern (Fig. 5C). The patterns were entirely visual in nature and required no reading literacy or numerical literacy beyond counting to 10. The test was not timed. Again the test was conducted on the same day as the recording but not simultaneously.

### EEG Recordings

Our recording paradigm involved a simple measurement of the resting EEG for 3 minutes when the subject was sitting still with their eyes closed. To carry out our recordings, we used the Emotiv EPOC wireless EEG headset with 14 gold plated electrodes (Sup. Fig. 1F) and 2 reference electrodes (M1, a ground reference point for measuring the voltage of the other sensors and M2, a feed-forward reference point for reducing electrical interference from external sources). The Emotiv EPOC is an inexpensive, portable and easy to use device, making it very advantageous for large-scale studies across multiple locations. In addition, it has a 12 hour battery life which is convenient for recording in remote locations where electricity may be absent or intermittent. Multiple studies have now shown that the EPOC is capable of measuring event-related potentials [35; 36; 37; 38; 39] as well as variation in EEG related to mood [40] and cognitive load [41]. Nonetheless, given its lower temporal resolution (128 Hz compared to ≥1kHz of clinical grade EEG) and higher electrode impedance (10-15 kΩ minimized by maintaining saline hydration of electrode sponges, compared to 2-5 kΩ of clinical grade electrodes) we sought to determine the suitability of the EPOC for our purpose by comparing the complexity measured in simultaneous recordings with the EPOC and a high end clinical grade EEG device. The very close results (Sup. Fig 3, described further down) motivated our use of the device.

Finally, we note that our eyes closed paradigm mitigated any artifacts from EMG signals arising from movement such as eye blink.

### Signal Complexity

We defined a measure of complexity in terms of the diversity of patterns represented in the waveform (Temporal Complexity or C_T_) as follows. For each channel or electrode we calculated the correlation *r_n_* of 100 non-overlapping randomly selected segments of *t* ms duration or *n* points where *n*= (*t*/1000)*sampling rate in Hz as defined below.

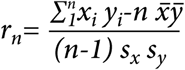
where 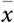 and 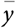 are the means of the signal and *s_x_* and *s_y_* are the standard deviation of the two segments compared (essentially the standard deviation of the amplitude distribution of the segment).

We then constructed the distribution of the (1−|*r*_*n*_|) ×100 values for each channel, which we call the *diversity distribution* (cumulative form shown in Sup. Fig. 2A) that provides a measure of how *different* the segments are. While the EEG from different electrodes in an individual could have a fair degree of variance, no particular region was distinct when averaged across all the sampled individuals (Sup. Fig. 2B). We then calculated the average diversity distribution across all channels and defined the complexity C_T_ for the individual as the median of the average diversity distribution.

By definition the measure could span a range from 0 to 100 where C_T_ = 0 would essentially be a highly repetitive pattern where at least half of the pairs of random segments had *r* = 1. At the extreme this would be a flat line. Conversely, a measure of 100 would reflect the maximal diversity where at least half of the segment pairs had a correlation of 0 such as would be expected with random noise, meaning the waveform rarely revisited the same pattern within the recording or analysis period.

We chose the period *t* = 750 *ms* for reasons described below corresponding to *n* = 96 points points. Measured thus the C_T_ varied from 35 to 96 across our sample, a 2.75-fold range (Fig 1A).

**Figure 1.**
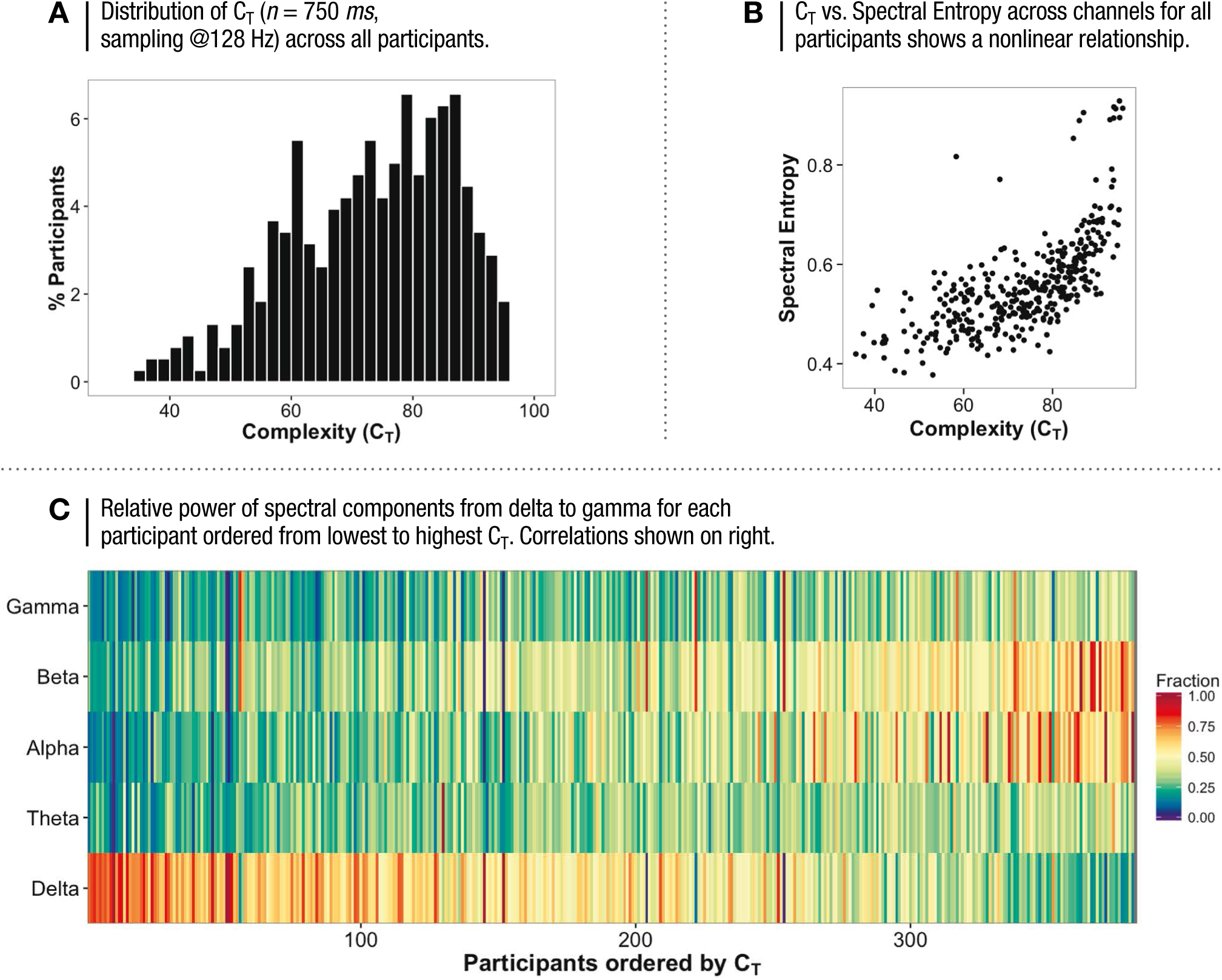
COMPLEXITY AND SPECTRAL PROPERTIES.

### Selection of segment duration (t)

A fundamental aspect of the measure is the length of segment compared. Shorter segments would have fewer possible configurations and therefore produce different results. Periods of synchrony with fine scale temporal structure have been found to vary in duration from 100 to 900 *ms* [42; 43]. We thus looked at complexity across our sample as a function of segment duration from 100 to 1250 *ms*.

At duration *t*=100 *ms* the mean C_T_ across our sample was ∼62, increasing sharply to ∼75 at *t* = 250 *ms* and then staying relatively flat, declining only slightly to 72 as the duration was increased to 1250 *ms* (Sup. Fig. 2C). However, as the duration increased from 250 to 750 *ms*, the coefficient of variation of C_T_ across our sample increased steeply from 10% to 18% at 750 *ms*, flattening out thereafter at 19% (Sup. Fig 2D; shown CV%). This is in sharp contrast to results obtained when the amplitude values of the signal were shuffled, destroying the temporal structure. In the shuffled signal C_T_ increased from 88 for *n*= 100 *ms* to 98+ for *n*=1250 *ms* indicating more repetitive temporal motifs in the signal relative to random. Correspondingly CV% decreased in the shuffled signal, tending towards 0 at 1250 *ms* from a max of 57% at 250 *ms* (Sup. Fig. 2D).

We thus chose the period *n* = 750 *ms* for analysis going forward as representative of the minimum duration at which complexity of the temporal sequence diverged maximally across the population.

The pattern of C_T_ versus sampling rate was also similar, increasing up to 32 Hz and then flattening out (Sup Fig. 2E) along with concomitant increases and flattening out of the CV% (Sup Fig. 2F) suggesting that most of the relevant signal had frequencies below 64 Hz (see next section for further discussion). Comparisons of the shuffled signal also increased with sampling rate with a similar pattern (but from 92 to 99+) while CV% again in stark contrast to the data, decreased, with increasing sampling rate, tending towards 0 as the sampling rate was increased.

### Effects of device properties and sampling resolution

C_T_ is fundamentally dependent on two factors of the measuring device, the electrode impedance and the sampling rate of the signal. Higher electrode impedance would add noise into the signal, particularly in the lower frequency ranges [44]. A lower sampling rate may lose important temporal structure that could distort C_T_. To determine the impact of these factors we compared recordings performed simultaneously on people fitted concurrently with both the EPOC and the clinical Neuroscan Version 4.3 (n=5, methods and data from Badcock et al, 2013) (Sup Fig 3A-D). Note that these simultaneous recordings were done for 13 minutes with eyes open and therefore represented a different experimental paradigm than our sample.

The EPOC signals were high-pass filtered with a 0.16 Hz cut-off, pre-amplified and low-pass filtered at an 83 Hz cut-off. The analog signals were then digitized at 2048 Hz and filtered using a 5th-order sinc notch filter (50 and 60 Hz) before being down-sampled to 128 Hz (company communication). The effective bandwidth of the signal is therefore between 0.16 and 45 Hz. In contrast, the Neuroscan samples at 1 kHz with band pass filtering between 1 and 100 Hz. To create as close a comparison as possible, and to remove 50 Hz noise, we applied 50 and 60 Hz notch filters to the Neuroscan (NS) signa.

The power spectrums of the simultaneously recorded EPOC and the NS signals were very similar when the NS signal was downsampled to 125 Hz by using every 8^th^ point in the recording (Sup. Fig. 3A). The EPOC had higher signals in the 0.16 − 1 Hz range that had been filtered out in the Neuroscan recordings. However, given that these are very low they are not likely to impact the complexity measure which involves changes in the signal on faster time scales. Moreover, any impact of this noise is likely to reduce the difference between groups and not artificially increase it.

We next looked at the impact of sampling frequency on C_T_. Much of the useful range of the EEG signal, given its spatial resolution and filtering of the signal by the skull, is likely to be within the 60-70 Hz range suggesting that the 128 Hz sampling of the EPOC would be sufficient. For instance, even rhythmic discharges of fast spiking bursts with spike rates greater than 100 Hz typically produce gamma range (30-70 Hz) oscillations in the LFP [45]. Nonetheless, high frequency activity in the EEG has been shown to relate to cortical processing [46]. Thus if large differences in waveform patterns exist between 45 Hz and 100 Hz or even beyond, C_T_ measured using EPOC could be a considerable underestimate. We therefore looked at C_T_ as a function of sampling frequencies of 25 to 500 Hz in the NS signal and 16 to 128 Hz in the EPOC.

Sup. Fig.3B shows the values of C_T_ for both devices averaged across the 5 simultaneous recordings as a function of sampling rate. C_T_ in both the EPOC and NS increased with sampling rate although C_T_ was marginally lower in the EPOC and flattened out beyond 64 Hz while C_T_ continued to increase slightly up to 250 Hz in the NS. Nonetheless at the resolution of 128 Hz the differences were small and systematic (Sup Fig. 3D). Thus, given the sampling constraints of the EPOC, it performed remarkably well.

We next looked at C_T_ in the EPOC and NS as a function of duration (at 128/125 Hz sampling for the EPOC and NS respectively). Here again the pattern was similar in the two devices (Sup. Fig. 3C), although C_T_ was marginally lower in the EPOC.

A comparison of the C_T_ values in the EPOC and the NS (duration 750 *ms*, sampling rate of 128/125 Hz) across the 5 individual is shown in Fig. 3D. C_T_ values in the NS were on average 2.3 points higher with a linear relationship (R^2^=0.62). Given the high similarity we concluded that the EPOC was an appropriate device for our study.

Finally we note that the C_T_ values in this sample of 5 people who were all graduate researchers at Macquarie University in Australia fell between 85 and 96, in the range that we found for people in our modern, educated and technology savvy group using the EPOC.

### Spectral Measures

To analyze the spectral properties of the signal we computed the component in each frequency band delta (0-4), theta (4-7.5), alpha (7.5-14), beta (14-30) and gamma (30+) for each channel as:

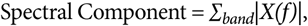
where *X*(*f*) is the Discrete Fourier Transform or DFT. We then calculated and reported the mean across all channels (Fig. 1C).

Spectral entropy was determined by computing the Shannon entropy of normalized power spectrum density (PSD_N_) as:

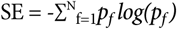
where *p*_*f*_ is the PSD_N_ for each frequency *f* and PSD = |X(*f*)|^2^. The reported value is the average across all channels (Fig. 1B).

### Statistical Tests

#### CONTROLLING FOR INTRA-PERSON VARIABILITY

To determine whether our results could be accounted for by intraperson variability rather than true population differences we measured resting, eyes closed EEG activity in 20 people in 10 separate sessions conducted over a week at different times of the day. We then used this to construct an average intra-person distribution and assess the probabilities of randomly picking groups with significantly different population means.

The coefficient of variation across channels within an individual, expressed as a percentage (CV%), had a median of 10%, compared to the CV% of 18% across our sample population (Sup Fig. 4A, n=400). People with higher complexity overall generally had lower CV% across channels, as low as 2%, while those with lower complexity overall were more likely to have greater cross channel variance (Sup Fig. 4B). This suggests that those with lower overall C_T_ have much more location-to-location variation and therefore potentially higher variability arising from differences in electrode placement.

We next looked at the variability within individuals (n=20) arising over separate recording sessions. The variation in C_T_ across sessions for each person was significantly narrower than the overall sample and also much narrower than the inter-channel variation (Sup Fig. 4B). Nonetheless, CV% within individuals ranged from 2% to 20% with an average of ∼10%. Furthermore, unlike the variation across channels, this variability was not correlated with the C_T_ value itself (Sup Fig. 4C filled circles) or with the inter-channel variation (Fig. 4D). This suggests that variation is more a consequence of state of mind than small differences in electrode placement.

We next constructed an average intra-person distribution by shifting each individual distribution such that the peak value was 0 (i.e. subtracting the peak value), and then averaging across them (Sup. Fig. 4E). Deviations from the peak represented the cross-trial variation in C_T_ points to the left and right of the peak in the average person. We then drew random samples of varying equal and unequal size (30 vs. 30, 30 vs. 60 which were typical of our size groupings in various comparisons) to determine the probability of a difference arising between the groups (Sup. Fig. 4F). The probability of a difference in the mean of ≥5 C_T_ points between two groups was less than 5% for both equal and unequal groupings. More significantly, the probability of choosing two groups with a difference in mean of 2.5 points or higher that was statistically significant at a 0.05 level by a *t*-test was 2×10^−3^ at the highest (Sup Fig. 4F inset). Thus it would be safe to assume that statistically significant results of 2.5 C_T_ points or higher obtained between groups of people along different experiential criteria would be highly unlikely to arise from artifacts of trial-to-trial variation within individuals.

#### STATISTICAL SIGNIFICANCE OF TRENDS

To determine the significance of the trends observed in Fig. 4, i.e. the shift in population means of C_T_ relative to income, education and other factors, we computed various statistics and probabilities. We first looked at the R^2^ of a best fit for income and education (logarithmic and linear respectively) as well as the correlation and their *p*-values, which provided an indication of the nature of the trend. We next computed the significance of an ANOVA (P_ANOVA_), which would provide the probability of a difference across the various bins. To determine the likelihood of such a trend appearing that could not be accounted for by intra-person fluctuations, we also computed the probability of finding a similar trend from shuffling the C_T_ values across the participants 1000 times including: (i) the probability of obtaining a significant ANOVA in the shuffled iterations that was ≤ 0.05 (*p*_*shuff1*_) or (ii) ≤ the *p*-value of the data (*p*_*shuff2*_) and (iii) the probability of finding a shuffled iteration with a trend that was positively correlated with p<0.05 to the data (*p*_*shuff3*_). These statistics are shown in Table 1.

### Principal Component Analysis

Principal Component Analysis was done using the FactomineR PCA function. Prior to application of the PCA function, all records without values in any of the columns were removed from the analysis (121/402 records). Secondly, given that the distributions of data across all variables except Travel and Education (i.e. Income, Fuel, Phone, Electricity, Population) were lognormal in nature, these variables were log transformed with a base 5 to create linear relationships among the variables. Individual component scores were calculated as the contribution of each element multiplied by the unscaled contextual factors.

## RESULTS

### A measure of complexity of the EEG signal

Complexity is not a clearly defined concept but in its broadest sense could be thought of as the diversity of possible configurations that could emerge from a nonlinear interdependent system. In the context of the brain, this could be the various patterns of activity produced by the communication of its billions of neurons. Much evidence now points to spatiotemporal patterns of neuronal activity as representative of memories and behaviors [47; 48; 49]. Furthermore these associations between spatiotemporal patterns of neuronal activity and memory and behavior have been found at various resolutions of measurement, from spiking patterns of individual neurons [48; 50; 51] to local field potentials (LFPs) measured with microelectrodes on the surface of brain tissue [52], ECoG [53; 54] and EEG [55; 56]. These field level potentials (LFP, ECoG, EEG) arise from the aggregate synaptic and spiking activity of the underlying neurons within the field of the electrode [43; 57]. The temporal structure or waveform is therefore grossly reflective of the underlying spatiotemporal patterns of neuronal activity and holds meaning about the internal state. It thus follows that the diversity of patterns of the measured waveform is likely to bear strong relation to the number of spatiotemporal patterns of underlying activity.

Complexity, when considered as diversity of the waveform structure, is very closely related to entropy. Typical measures of EEG signal entropy involve its decomposition into either its spectral or wavelet components followed by a subsequent analysis of the diversity of these components, and have been found to have relationships to both depth of anesthesia and disease states [58; 59; 60]. Motivated by studies that demonstrate stimulus related changes in the structure and synchrony of waveforms in short epochs [43] and the loss-less transmission of complex waveforms that are related to behavior [42; 53], our measure is related to these measures of entropy, but is constructed on the premise that it is the whole waveform structure that is important (and therefore the interspersion of spectral and wavelet components). One might imagine the analogy to a line of text where the relative positioning and spacing of letters is what confers meaning as opposed to how many of each letter there are. Our measure thus maintains the signal in its intact form, identifying the diversity of complex waveform patterns in the signal. A complete description of this measure of complexity and the factors considered in its definition are described in the methods, along with supplementary figures. By definition our measure could span a range from 0 to 100 where a flat line would be 0 and both white noise and a structured highly non-repetitive pattern over our measurement would be 100. Measured in this way, complexity spanned a ∼2.75-fold range across our sample with values from 35 to 96 (Fig. 1A).

### Relation of C_T_ to measures of Spectral Properties and Entropy

The greater the number of distinct waveform patterns in the system, the greater the complexity. To create a larger number of waveform patterns however, one must necessarily have a wide range of frequencies with various phase relationships. We therefore looked at the relationship between our complexity measure to the relative power in the different frequency ranges delta (1-4 Hz), theta (4-7.5 Hz), alpha (7.5-14 Hz), beta (14-30 Hz) and gamma (30+ Hz). As would be expected, we found that higher temporal complexity was generally associated with a higher fraction of frequencies above 4 Hz, although none of the higher frequency bands were more specifically related to the measure (R_2_ of linear fit 0.19 for both theta and gamma, 0.41 for beta and 0.44 for alpha). Conversely, this was accompanied by a concomitant decrease in the delta power (R^2^ of linear fit 0.54). This pattern is easily visualized in Fig. 1C showing the summed power in each frequency band for each participant ordered by complexity.

We then compared the spectral entropy of the signal (the Shannon entropy of the power spectrum, see methods) to our measure of complexity (Fig. 1B). The two measures were positively correlated (*r*=0.66) but had a nonlinear relationship, an indication that the measures were related but yet carried different information. For instance, people with low spectral entropy values of 0.45 could have complexity values ranging from 40 to 80 indicating that the diversity of patterns constructed from the same spectral composition varied substantially.

### Complexity, Socioeconomics and Modern Experience

We next explored whether the complexity of the EEG reflects the divergent complexity of human experience in the transition from pre-modern to modern society. To do so we compared the complexity of different groupings of income and experience.

Before describing these results, however, we note that our study involves comparison of the complexity of the resting EEG signal from just a 3-minute snapshot across a large number of people. It is thus possible that the variation in C_T_ across the population was simply a function of intra-person variability arising from differential placement of electrodes or state of mind at the time of recording. We therefore examined the variability arising within 20 individuals across 10 sessions conducted at various times over a one week period to determine levels of statistical significance in differences among random groups drawn from the average intraperson distribution (described in detail in Methods, Sup. Fig 4).

The probability of a difference in the mean between two groups being ≥5 C_T_ points was less than 5% for both equal and unequal groupings of sizes similar to our experience groupings. More significantly, the probability of choosing groups with a difference in mean of 2.5 points or higher that was statistically significant at a 0.05 level by a t-test was 2×10^−3^ at the highest (Sup Fig. 4F inset). Thus it would be safe to assume that statistically significant results of 2.5 C_T_ points or higher obtained between groups of people along different experiential criteria would be highly unlikely to arise as a consequence of trial-to-trial variation within individuals.

We first compared gender and age groupings. Our sampling was such that each income group in our sample was equally distributed across males and females and decadal age groups, thus comparisons along these dimensions would not be likely to reflect second order differences along other dimensions.

The distribution of EEG complexity of males and females were not significantly different (Fig. 2A) with mean ± SEM for males at 74.07±0.96 and Females at 72.11±0.98 (p>0.15 both KS test and *t*-test).

**Figure 2.**
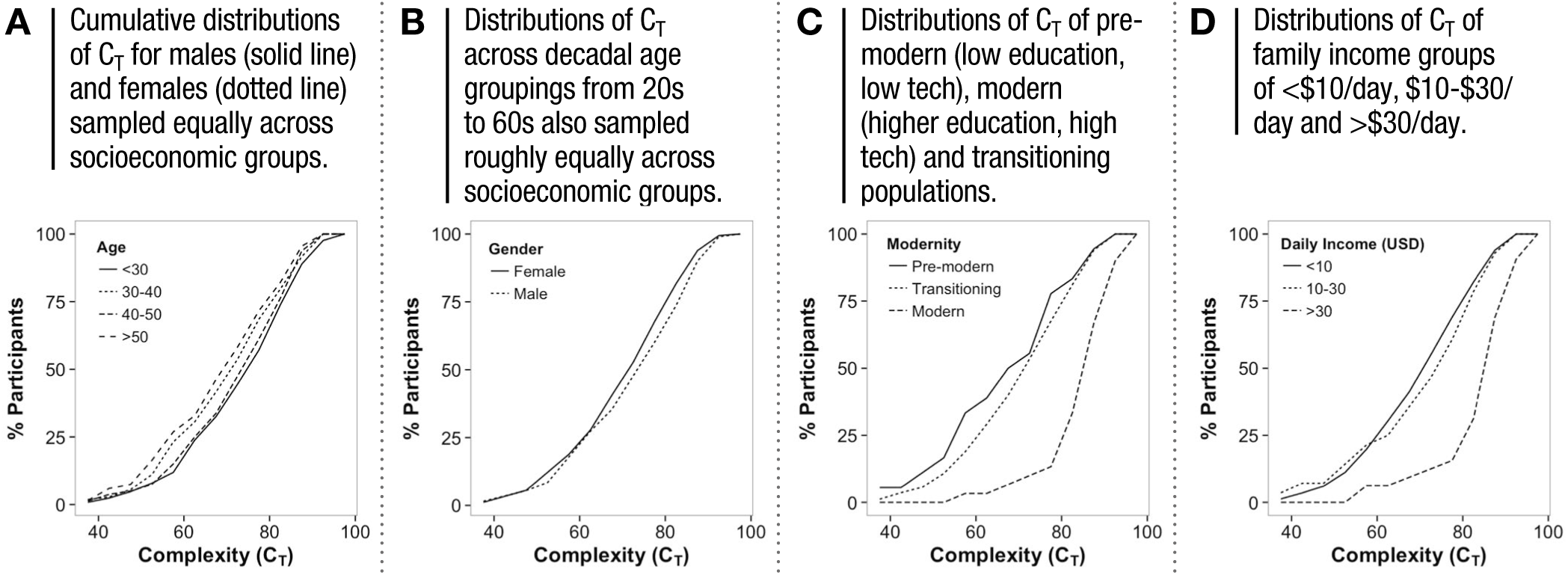
DISTRIBUTIONS OF COMPLEXITY FOR AGE, GENDER, MODERNITY AND INCOME.

Similarly there was no significant difference between any successive decadal age groups (Fig. 2B; mean ± SD *age 21-30:* 75.09±1.16, *30-40:* 71.99±1.31, *40-50:* 73.97±1.45, *50-60:* 70.35±1.61, 0.18<p<0.6 by KS test, ANOVA *F* = 2.23 *p* = 0.08 old), although the 21-30 age group was significantly higher than those 51-60 (*p*=0.02).

We next divided the population into three groups along a dimension of modernity (Fig. 2C): (i) Pre-modern, referring to those with primary education or below and limited use of technology (no mobile phone, internet or owned vehicles), (ii) modern, referring to college educated with extensive use of mobile phones, internet access and owned vehicles, and (iii) transitioning, referring to those at various stages of education and technology use between these two extremes. Complexity of the pre-modern, transitioning and modern groups were (mean±SEM) 71.53±2.65, 72.07 ±0.73 and 86.23±1.53 respectively. Pre-modern and transitioning populations were not significantly different (*p*=0.97 by KS test). In contrast both were dramatically distinct from the modern group (*p*= 1.8 × 10-5 *t*-test, 6.1 × 10-9 by KS test). The dramatic shift of the modern group suggests a distinctly non-linear jump in complexity at a certain threshold of modernity.

Given that much of modern experience is enabled by income, we next looked at the difference in C_T_ across different income groups (Fig. 2D). We found a similar pattern whereby the population with family income below $10/day and between $10 and $30/day were largely similar with means±SEM of 71.82±0.74 and 72.9±2.82 respectively (*p*=0.37). In contrast, the group with incomes of >$30/ day had mean±SEM of 85.28±1.68, dramatically different from both other groups (*p*=3.5 × 10-10 *t*-test, 1.8 × 10-5 KS test), once again suggesting that the increase in complexity is not linear.

To probe in more depth the factors that contributed to this difference we binned participants more finely along various income and experience dimensions (described in Sup. Table 1) and looked for systematic trends in the population complexity (shown are mean ± SEM of C_T_)(Fig. 3). However given that these were trends across a number of bins, for further assessment of their significance and likelihood that they were a consequence of intra-person variability we computed the significance of an ANOVA across these bins (*p*_ANOVA_) and compared it to the probability of finding a similar trend from shuffling the C_T_ values across the participants 1000 times including the probability of obtaining (i) a significance level in the ANOVA that was less than 0.05 (*p*_*shuff1*_), (ii) a significance level in the ANOVA that was less than the *p*-value of the data (*p*_*shuff2*_) and (iii) the probability of finding a shuffled iteration that was positively correlated to the trend in the data (*p*_*shuff3*_) (detailed in Methods). While differences between specific groups and the ANOVA are shown in the text with standard statistics between groups, the comprehensive trend statistics are shown only in Table 1.

**Figure 3.**
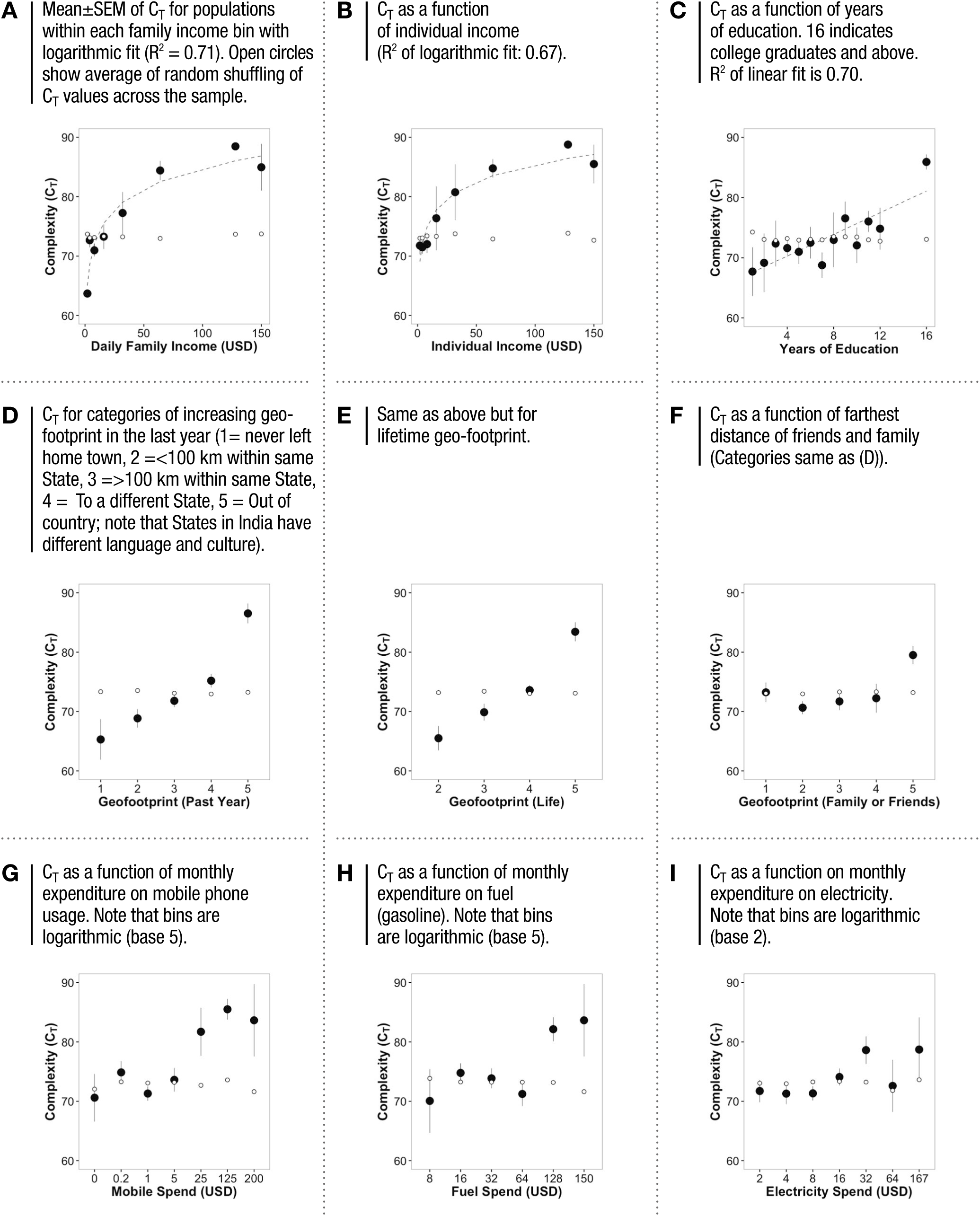
SCALING OF COMPLEXITY WITH VARIOUS EXPERIENCE FACTORS.

**Table 1.**

| No. | Life Context Factor | Figure | Range (Max -Min) | P-value (Max, Min) | P-shuff1 | P-shuff2 | P-shuff3 |
| --- | --- | --- | --- | --- | --- | --- | --- |
| 1 | Age | 2A | 5 | 2.40E-02 | 6.2E-02 | 9.6E-02 | 3.1E-02 |
| 2 | Gender | 2B | 3 | 1.60E-01 |  |  |  |
| 3 | Modernity | 2C | 15 | 1.80E-05 | 5.8E-02 | 0.0E+00 | 3.4E-02 |
| 4 | Daily Family Income (USD) | 2D | 13 | 3.50E-07 |  |  |  |
| 5 | Daily Family Income (USD) | 3A | 21 | 7.02E-03 | 5.8E-02 | 0.0E+00 | 5.1E-02 |
| 6 | Daily Individual Income (USD) | 3B | 14 | 5.93E-03 | 5.8E-02 | 0.0E+00 | 2.0E-02 |
| 7 | Years of Education | 3C | 18 | 8.54E-04 | 5.1E-02 | 0.0E+00 | 0.0E+00 |
| 8 | Geofootprint Past Year | 3D | 21 | 6.80E-06 | 3.8E-02 | 0.0E+00 | 4.8E-02 |
| 9 | Geofootprint Life | 3E | 18 | 4.31E-09 | 3.8E-02 | 0.0E+00 | 4.8E-02 |
| 10 | Geofootprint Family/Friends | 3F | 6 | 6.31E-03 | 6.2E-02 | 0.0E+00 | 1.0E-02 |
| 11 | Mobile Phone Spend | 3G | 11 | 1.68E-05 | 6.6E-02 | 0.0E+00 | 1.0E-01 |
| 12 | Fuel Spend | 3H | 14 | 1.49E-01 | 6.8E-02 | 1.0E-02 | 3.7E-02 |
| 13 | Electricity Spend | 3I | 7 | 2.68E-01 | 5.5E-02 | 7.9E-02 | 3.1E-02 |
| 14 | Name Task Score | 5A | 19 | 6.34E-05 | 4.1E-02 | 4.1E-02 | 5.5E-02 |
| 15 | Pattern Task Score | 5B | 29 | 5.14E-04 | 3.1E-02 | 0.0E+00 | 1.4E-02 |

#### INCOME

Consistent with the nonlinear jump in C_T_ in Fig 2D, the relationship between C_T_ and income was best fit by a logarithmic function (Fig. 3A,B R^2^ = 0.71 for Family Income and 0.67 for Self Income, ANOVA *F* = 6.08 *p* = 9.9 × 10^−7^ family income and *F*= 5.88 *p* =1.9 × 10^−6^ individual income), indicating that the population C_T_ shifted sharply to the right with increasing family income up to ∼$30/day and increasing individual income up to about ∼$17/day, leveling out thereafter. The difference in complexity between the lowest and highest income groups was 21 points. We note however that between the range of $3 and $10 where the largest fraction of our sample (and humanity) resides, there was no significant difference.

#### EDUCATION

Education represents the most formalized access to knowledge of increasing complexity. Indeed mean C_T_ of the population scaled systematically with years of education and could be best fit by a linear function (Fig. 3C, R^2^ = 0.70, ANOVA *F* = 4.55 *p* = 8.1×10^−7^) with an 18-point difference in the mean between unschooled and college-educated groups (16 years). We note that those who drop out of school at various stages without going to college, tend to have gone to lower income government schools where standards of education are relatively poor, while the college goers have also had the benefit of a better private schooling. This could be a major reason for the greater variance and the sharper increase from 12-16 (college) years relative to 1-12 years of education.

#### GEOFOOTPRINT

Geofootprint refers to the distance a person has traveled in the world or the geographical space of exploration. Travel is highly dependent on access to modern transport and requires a wide repertoire of ability from planning to spatial navigation to navigating new languages and cultures. This measure thus stands as gross proxy for an integrated range of different stimuli that a person is likely to have encountered in their lifetime. To get an approximate measure of a person’s geofootprint we classified their travel into six categories reflecting the distance and complexity of their travel. Category 1 referred to no travel outside of ones hometown, 2 referred to trips to places that were less than 100 km away and within the same State and were typically a day trip, Category 3 referred to trips to places that were within the same State but more than 100 km away and therefore required overnight stay and more planning. Category 4 was to another State, and 5 was out of the country. Crossing State borders in India represents a transition into a different language and culture and therefore adds a significant layer of complexity. We then identified the category that defined the maximum extent of their travel in the last year as well as in their lifetime. We also identified the category that defined the maximum geographic range of their social network (Farthest friends and family).

Mean C_T_ scaled systematically by 3 or 4 points for each successive category of geofootprint (Fig. 3D,E, ANOVA *F* = 11.35 *p* = 9.1 × 10^−9^ past year, *F* = 13.27, *p* = 2.9 × 10^−8^ life). Populations with overseas footprints within the last year however had C_T_ values 8 point higher than those who had not left the country, while populations with overseas footprints over their entire lifetime had C_T_ values 6 points higher than those who had never left the country. Overall, the difference in C_T_ between populations who had never left their hometown to those who had traveled overseas in the last year or their lifetime was 21 and 18 points respectively. In contrast, there was a far less significant difference between populations with geographically wider social networks with only category 5 (overseas family or friends) having any significance (Fig. 3F, ANOVA *F*= 5.55 *p* = 2.4 × 10^−4^).

#### COMMUNICATION TECHNOLOGY

We also looked at mobile phone and Internet usage. Since all users of the Internet in our sample were college educated and had similar hours of usage, we were unable to make any useful comparison. This is therefore excluded. However we found that populations with increasing mobile phone usage were not substantially different until the level of usage went beyond 300 Rs. (∼$5) per month (equivalent to ∼ 300 minutes of talk time or 1.25GB of data transfer per month) at which point there was a dramatic jump in C_T_ of 8 points (Fig. 3G, ANOVA *F*= 4.65, *p* = 1.6 × 10^−4^). Note the logarithmic trend, similar to income.

#### FUEL CONSUMPTION

Fuel stands as a proxy for daily activity and local travel. Here again we found no significant change in C_T_ as a function of increasing fuel consumption until fuel consumption was $128/month or more (Fig 3H, ANOVA *F* = 3.78 *p* = 0.003) where C_T_ was 8 points higher than all values at lower fuel consumption. Overall the effects were significantly less than for Geofootprint, income and education.

#### ELECTRICITY CONSUMPTION

Finally, we looked at C_T_ as a function of electricity consumption. There was some evidence of a threshold dependent trend as seen for mobile spend with a greater likelihood of an increase when electricity consumption exceeded ∼$32/month, however the effects here were not significant (ANOVA *F*= 1.94 *p* = 0.071).

Overall, the statistics were highly significant (Table 1) with very small probabilities that these patterns could have emerged simply from intra-person or sampling variations of the populations. Finally we note that both the related measures of spectral entropy and delta fraction had directional similarities but without the consistent statistical significance of this measure (not shown).

#### PARSING THE CONTRIBUTIONS OF INDIVIDUAL FACTORS

Taken together our results suggest that the complexity of the EEG reflects a host of economic and experiential factors, most significantly income and travel. Income serves as a universal enabler for the experiential factors of education, travel, communication and energy consumption. People with higher income therefore have greater access to all elements, and all factors would likely strongly co-vary with income. Thus while there were clear shifts in the mean population C_T_ as a function of most experiential factors, it is not clear how much each factor contributes independent of the others. Indeed all of the factors we considered were positively correlated with one another, with income most tightly correlated with education and mobile phone use (Fig. 4A). In contrast, geofootprint (travel year and life) had the weakest correlation with income, education and other factors. The systematic shift in the population mean of C_T_ with geofootprint was evident even when separated by income (Fig. 4B) or education (Fig. 4C) categories, and was apparent even for the high income and education groups although they spanned only two of the travel categories (geofootprint 5 or 6).

We next performed a principal component analysis to identify any further dimensions of experience that could be disaggregated from one another, and income in particular (see Methods for details).

**Figure 4.**
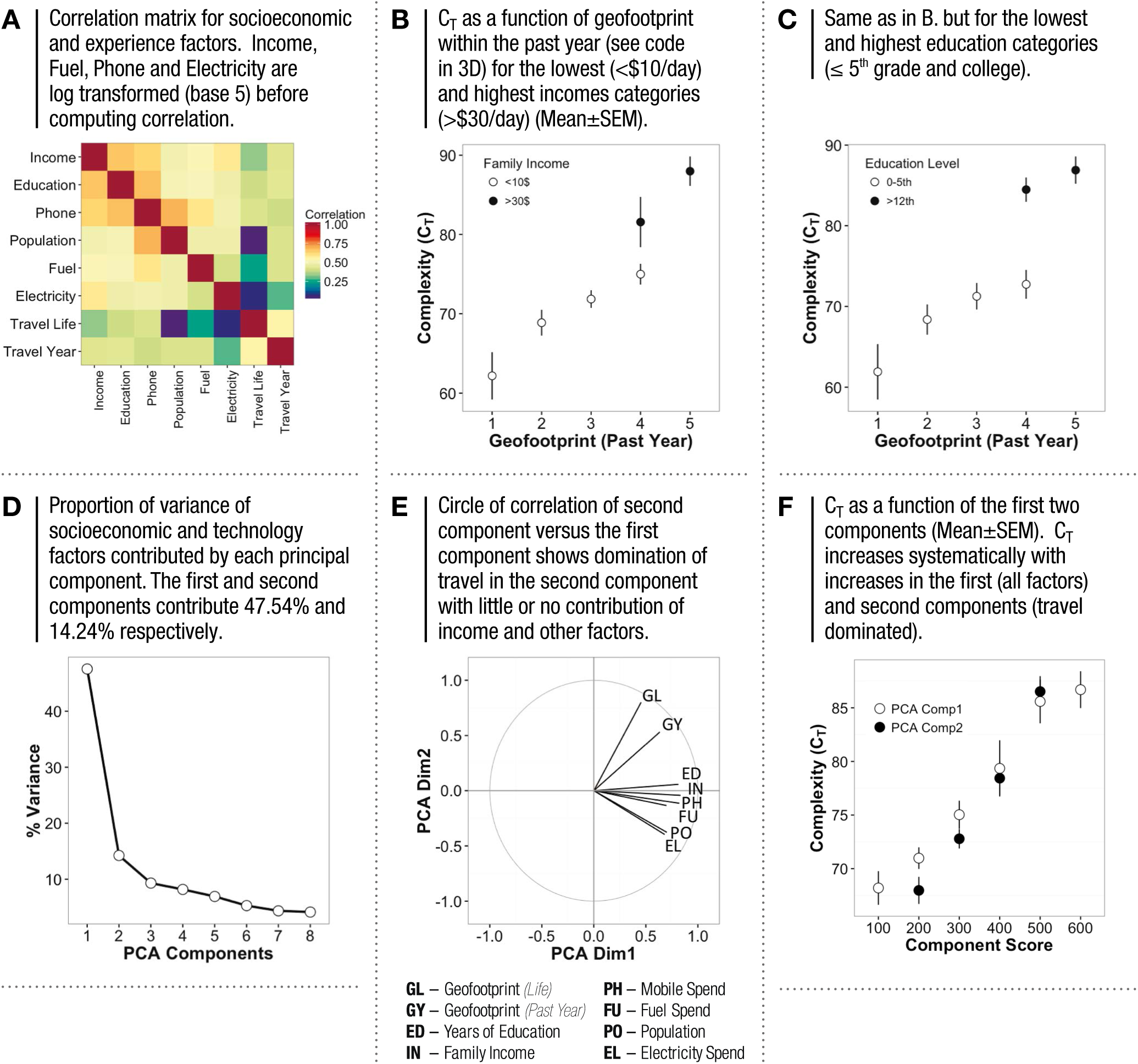
PARSING OUT EFFECTS OF EXPERIENCE FACTORS FROM INCOME.

The first principal component (Comp1), where all factors of experience essentially co-varied with income, accounted for approximately 47.54% of the variance in C_T_ (Fig. 4D). The second component (Comp2) contributed 14.24% of the variance and was dominated by travel (geofootprint), while all other factors made little to no positive contribution (Fig. 4E). Thus travel appears to be an experiential choice that is more individual and independent relative to all other factors. We next looked at C_T_ as a function of individual scores constructed for each of these components (Fig. 4F). Indeed C_T_ scaled systematically as a function of both components (R^2^=0.94, P_ANOVA_ *p* = 2.3 × 10^−9^ (Comp1), R^2^ = 0.96, P_ANOVA_ 4.6 × 10^−10^ (Comp2)).

## COMPLEXITY AND FUNCTIONAL CAPACITY

A major question that arises when attempting to interpret these results is the relationship between this measure of complexity (C_T_) and the functional capacity of the individual. While this warrants in depth investigation with respect to various dimensions of cognition, we made a rudimentary first attempt to quantify two functional aspects, namely the speed of free recall of names of people or places and the ability to recognize patterns (Fig. 5). These tests were administered on the same day as the EEG recording but were not simultaneous with the recording, which required the participant to be still with their eyes closed.

**Figure 5.**
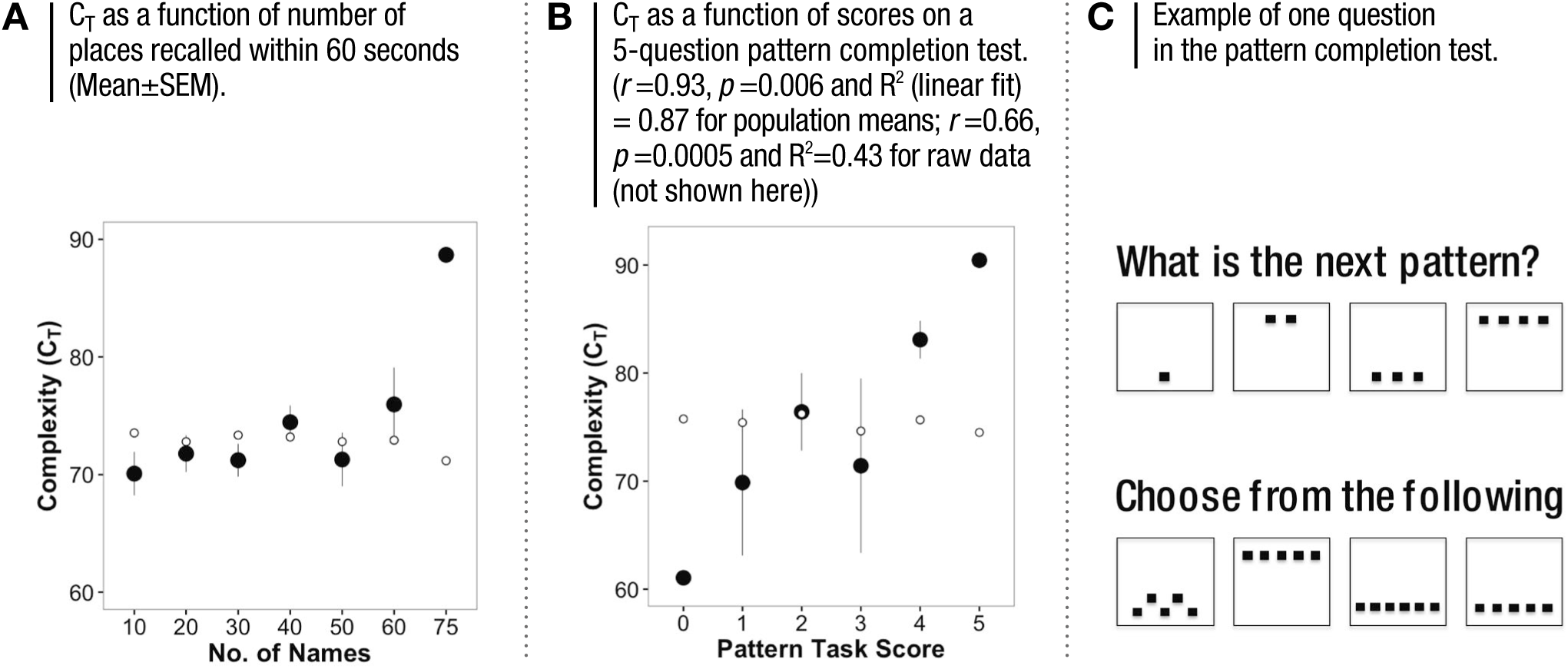
COMPLEXITY AND COGNITIVE TESTS.

### NAME RECALL

This test, which we administered only to people with writing literacy (n = 346) required writing as many names of people and places as possible within a one minute period. We note that this could reflect a larger pool of names available for recall, a faster rate of retrieval and/or greater motor facility and is a coarse test at best.

Here we found that C_T_ increased only slightly as the number of names recalled increased from 10 to 50 but then grew dramatically higher beyond (Fig. 5A). The overall trend was not significant (ANOVA *p* = 0.12), however the group with scores >60 had an average C_T_ value 12 points higher than the 50-60 names group (*p* = 6.3 × 10^−5^) and 19 points higher than the 0-10 group. The nonlinear trend hints that higher C_T_ may be associated with non-linear gains in this cognitive function. However, the relatively low correlation of the raw scores to complexity (*r* = 0.12 *p* = 0.018) suggests that this measure is at best an indirect measure of recall or retrieval speed.

### PATTERN COMPLETION

The pattern completion test used a modified and abridged version of the Raven’s progressive matrices and consisted of five multiple choice questions of increasing difficulty, a sample of which is shown in Fig. 5C. The test was administered to 28 people across two locations (one urban and one rural). The patterns were visual in nature and required no formal verbal or numerical literacy beyond counting to 10. Unlike name recall, we found that EEG complexity scaled linearly with test scores. The correlation between complexity and raw scores of all 28 people (not shown) was 0.66, (*p_r_* = 1.5 × 10^−4^), while the shift in mean complexity within each score group (Fig. 5B) followed a linear trend with a correlation of 0.93 (*p^r^* = 0.006, p_ANOVA_ = 0.007 and R^2^ of linear fit of 0.43). We note that this was a far stronger relationship compared to the related measure of spectral entropy where the correlation was 0.4 (*p^r^* = 0.03).

All together this suggests that our measure of complexity strongly reflects both experience and cognitive function.

## DISCUSSION

The brain lies at the core of our sense of what is human. Most of the understanding we have built thus far, however, comes from work done in the developed world, and with subjects that are, most often, university students representing only a very small privileged fraction of humanity. The implicit assumption is that findings from these studies extend easily to all of humanity.

This study is the first of its kind, extending far beyond the populations that reside around universities and into populations relatively untouched by modern advances, with four important results. First, the complexity of the EEG signal varies severalfold across individuals. Second, the complexity of the signal is nonlinearly related to the increasing complexity of modern living as well as to income. The relationship with income is logarithmic in nature similar to that reported between income and brain surface area for a study conducted in the United States [30]. Given the structure of this relationship parity in complexity was roughly achieved at a family income of ∼$30/day or individual income of $17/day. Third, geofootprint or travel of the individual has the most dramatic independent contribution to complexity relative to all other factors including education. Fourth, complexity is strongly correlated with scores on a pattern completion task (and weakly with memory retrieval) indicating that higher complexity implies clear cognitive gains. Taken together, these results demonstrate a wide range in the dynamical complexity of the human brain that is profoundly related to socioeconomic context and geofootprint and has cognitive relevance.

We note poignantly that 80% of the world’s population falls below the threshold of $30/day of family income beyond which EEG complexity distributions show parity across incomes.

A crucial question that arises is therefore the direction of causality – does higher complexity confer higher income or does higher income confer higher complexity? Does travel increase complexity or does higher complexity spur greater travel and exploration?

The malleability of complexity and functional capability with different dimensions of experience is a critical question that must be addressed going forward, with significant implications for how we might approach a global development agenda. Indeed, although it was once thought that the adult human cortex was not capable of substantial rewiring, there has been mounting evidence to the contrary [10; 61; 62; 63]. Thus while a greater ability to make sense of new situations and patterns may propel greater geo exploration, such exploration will likely in turn enhance complexity, leading to meaningful cognitive benefits. In this construct interventions that lower barriers to physical mobility may deliver the most meaningful gains by allowing people to explore a wider range of environments, and warrants deeper investigation.

Finally our study warns against extending results obtained from small or socioeconomically limited groups to global populations. We also note that our study demonstrates that, with new cost-effective technologies available, it is possible to take neuroscience investigation out of the lab and to a much larger global population.

## ACKNOWLEDGEMENTS

A very special thanks to Sathish S, Aravind and Govind for managing the field EEG recordings and surveys, as well as S. Ravi Shankar who enabled smooth coordination of the process.

We also thank the various staff at Madura Microfinance who facilitated access to the villages and recruitment of participants, SciSphere, India for providing us survey tools, access to demographic and economic data for location selection and data support, and Nicholas Badcock and Betty Mousikou at Macquarie University in Sydney, Australia for providing us with the simultaneously recorded Emotiv EPOC and Neuroscan data.

---

**Sup. Figure 1.**
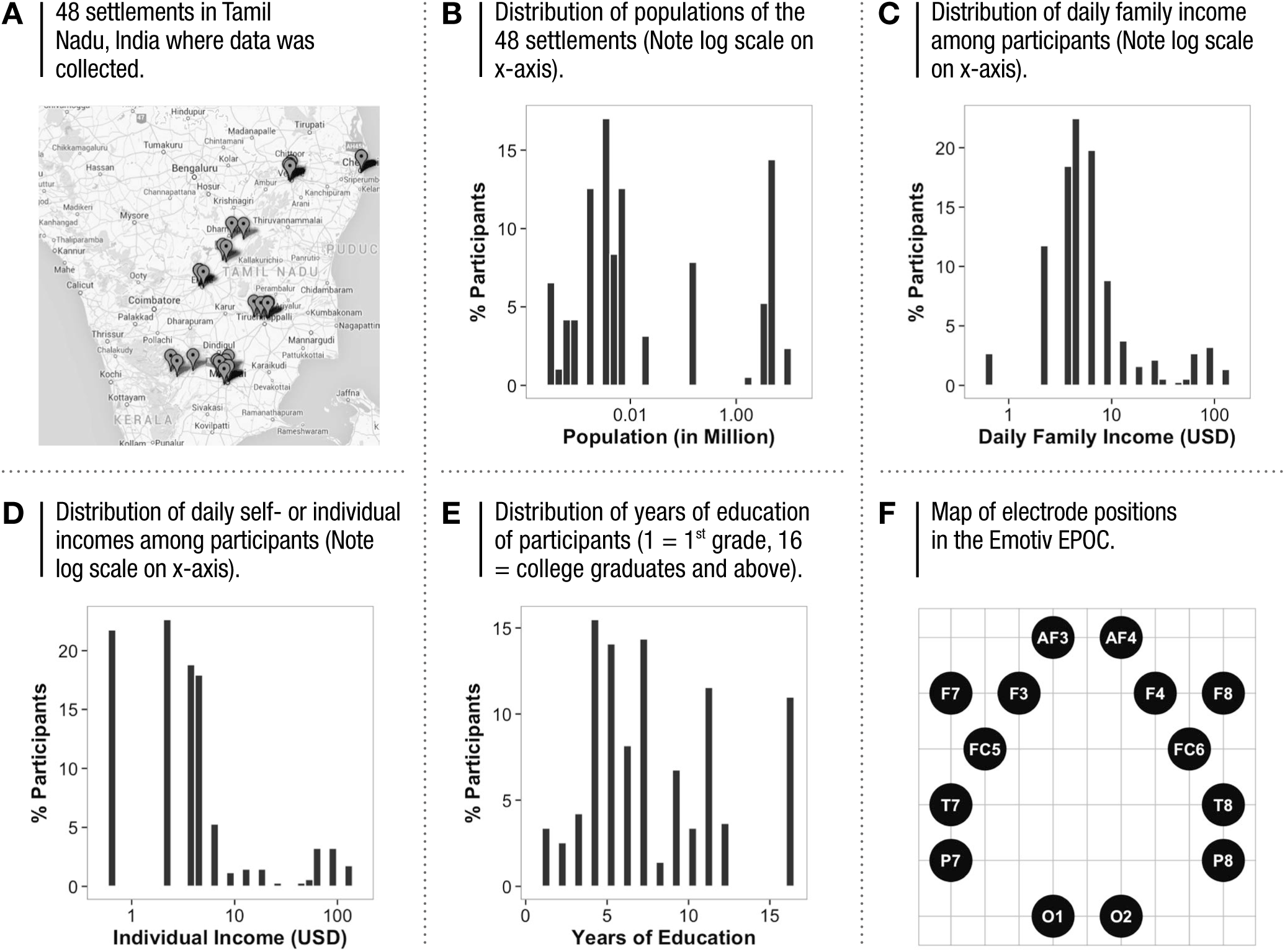
SAMPLING ACROSS A BROAD RANGE OF HUMANITY.

**Sup. Figure 2.**
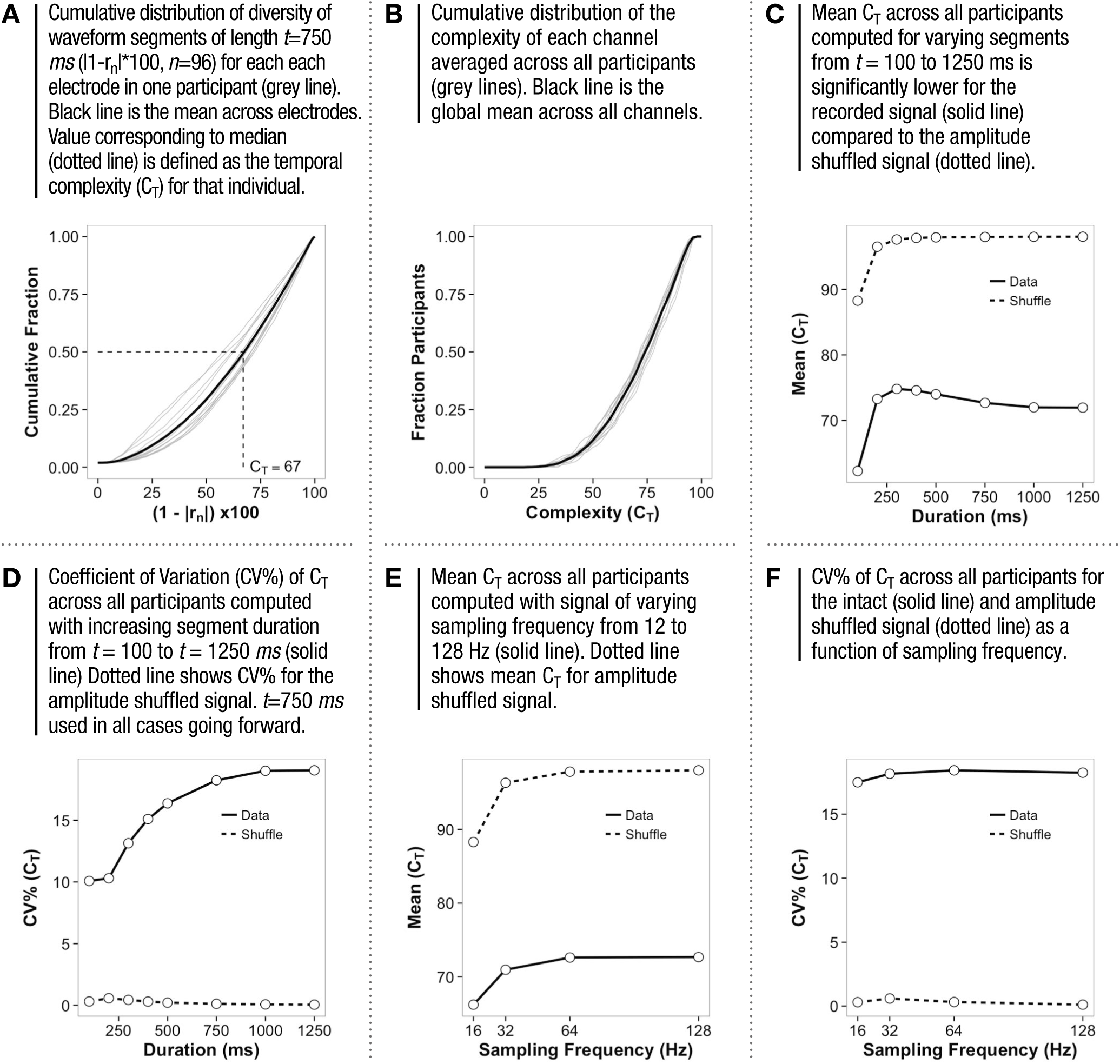
DEFINING A MEASURE OF COMPLEXITY.

**Sup. Figure 3.**
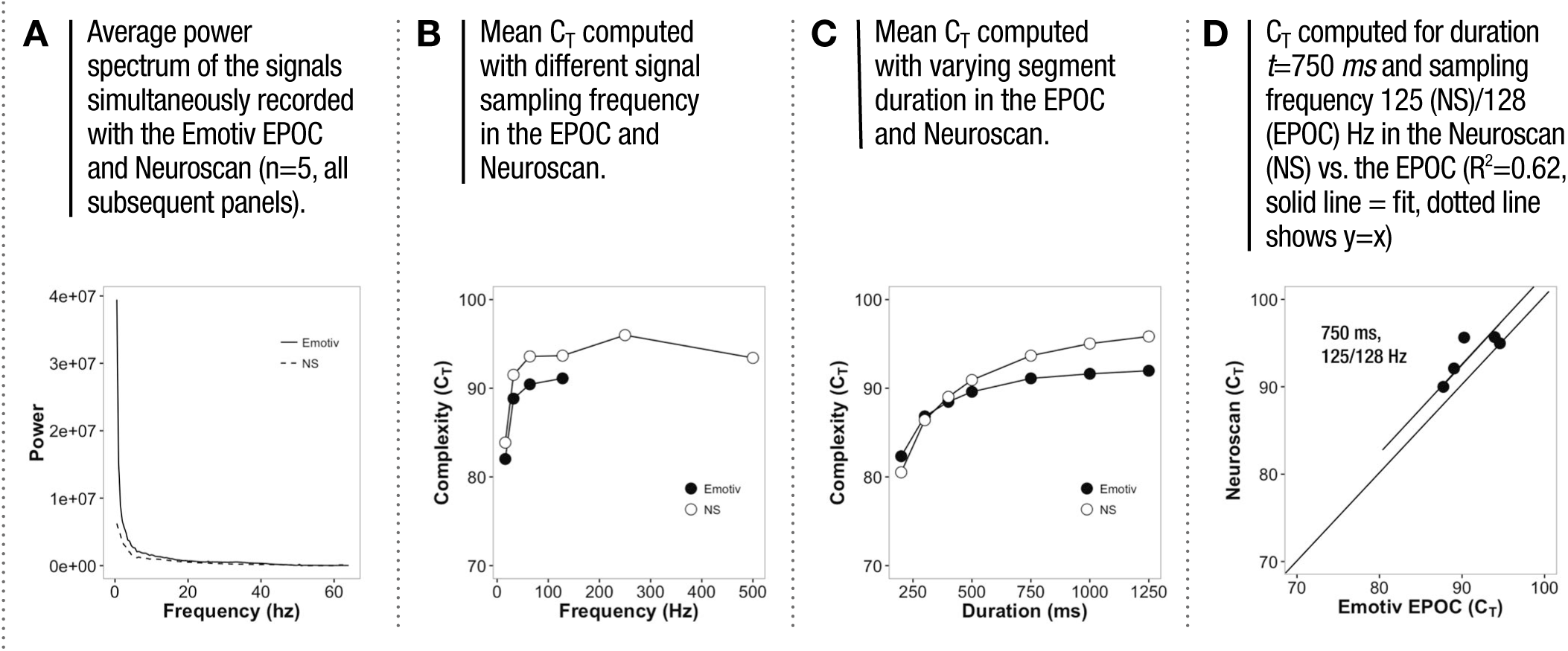
COMPARISON OF THE EMOTIV EPOC TO NEUROSCAN.

**Sup. Figure 4.**
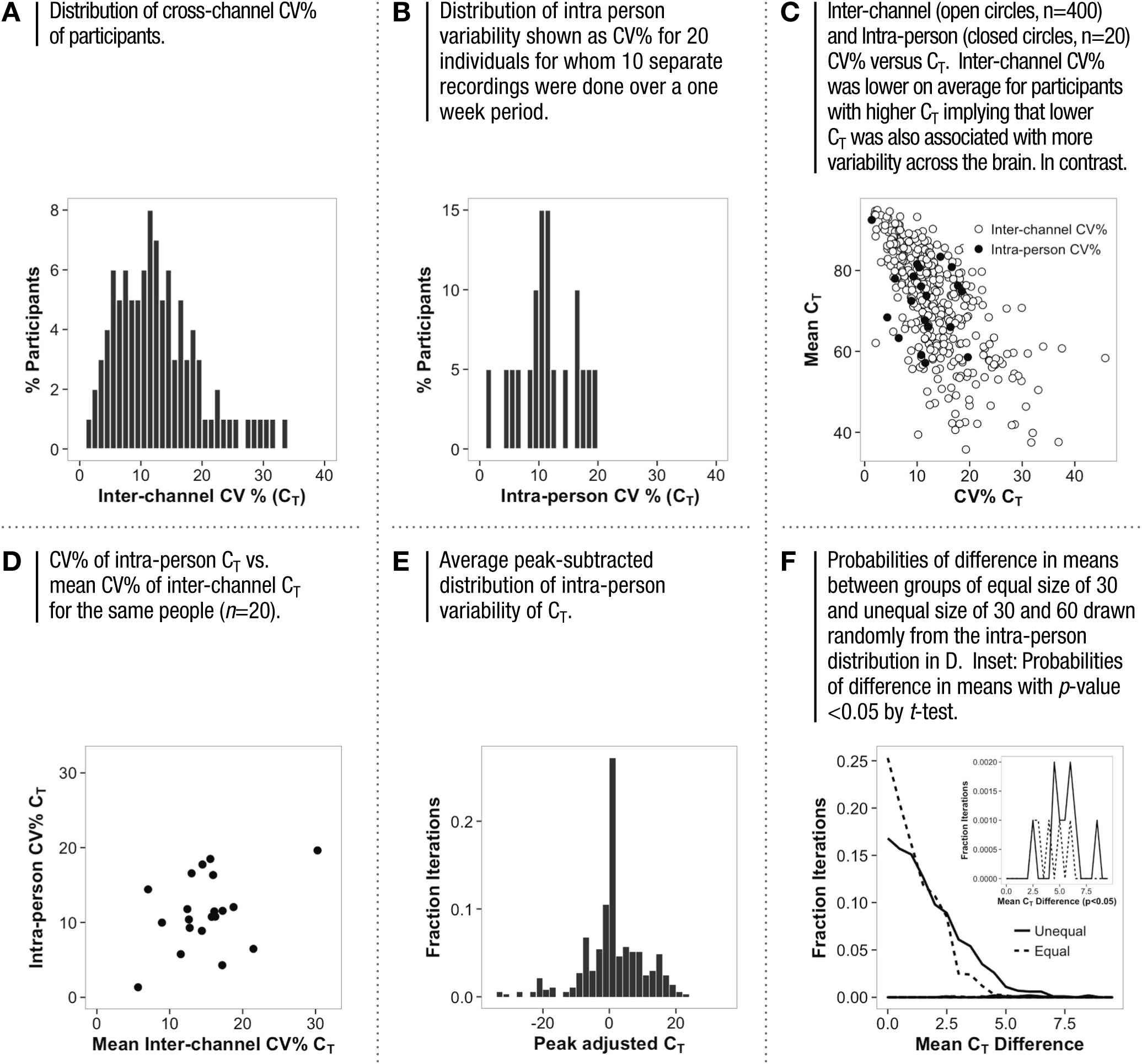
INTRA-PERSON VARIABILITY OF C_T_.

**Sup. Table 1.**

| No. | Feature | Description |
| --- | --- | --- |
| 1 | Family Income | Monthly income in Indian Rupees of the total household income. 60 Rs = \$1. 50,000 Rs. is ~\$850 or \$28 per day. |
| 2 | Individual Income | The individual's self-earned monthly income in Indian Rupees. |
| 3 | Education Level | Number of years of completed education. Of those who attended college, all in the sample completed it. Note all college grads are marked as 16 years of education, although some attended 3 year college. |
| 4 | Geofootprint (Past year) | This indicates the distance from home they have traveled in the one year before the date of survey. The question was asked in the form of the following categories:<br>1: Within home town<br>2: Within 100 km from hometown in the same State (typically a day trip)<br>3: >100 km from hometown in the Same State (typically requires overnight stay)<br>4: Within 100 km from hometown to a different State (note that State boundaries in India are based on language so different State indicates a different language)<br>5: >100 km from hometown to a different State<br>6: To a different country |
| 5 | Geofootprint (Lifetime) | This indicates the distance from home they have ever traveled in their life for any reason at any age. The categories are the same as above. |
| 6 | Farthest Friends or Family | This indicates the furthest distance from home that they have either friends or family, essentially an indication of the geographic distance of their social network. The categories are once again the same as above. |
| 7 | Fuel Usage | This refers to the amount they spent on purchasing petrol or gasoline in the previous full month shown in Indian Rupees. The price of petrol was ~ Rs. 75/liter or \$4.75/ gallon at the time of survey. The range of fuel purchase ranged from 0 (no vehicle) or going from 1 liter all the way to 160 liters or 42 gallons, translating to miles traveled in the month of anywhere from 20 to 1500. |
| 8 | Electricity Usage | This refers to the amount they spent on household electricity in the previous full month shown in Indian Rupees. The price of electricity at the time of survey ranged from Rs. 1 to Rs. 4 per unit or kWh. The tariff was a sliding scale with lower prices for those consuming lower levels of electricity. Consequently an exact conversion to units consumed was not readily possible. |
| 9 | Mobile Phone Usage | This refers to the amount spent on mobile phone communication in the previous full month shown in Indian Rupees. |
| 10 | Internet Usage | Here we asked for hours of Internet use. However, since it was used only by highly people with income >50,000 Rs. per month, the data was insufficient to make relevant comparisons. |

## REFERENCES

[1] I. Ortiz, and M. Cummins, Global Inequality: Beyond the Bottom Billion: A Rapid Review of Income Distribution in 141 Countries. Unicef Social and Economic Policy Working Paper (2011).

[2] C. Lakner, and B. Milanovic, Global Income Distribution: From the Fall of the Berlin Wall to the Great Recession. The World Bank Policy Research Working Paper 6719 (2013).

[3] P. Medini, Experience-dependent plasticity of visual cortical microcircuits. Neuroscience 278 (2014) 367–84.

[4] J.W. Schnupp, and O. Kacelnik, Cortical plasticity: learning from cortical reorganisation. Curr Biol 12 (2002) R144–6.

[5] M. Sur, J. Schummers, and V. Dragoi, Cortical plasticity: time for a change. Curr Biol 12 (2002) R168–70.

[6] C. Xerri, Plasticity of cortical maps: multiple triggers for adaptive reorganization following brain damage and spinal cord injury. Neuroscientist 18 (2012) 133–48.

[7] S. Chaudhury, V. Sharma, V. Kumar, T.C. Nag, and S. Wadhwa, Activity-dependent synaptic plasticity modulates the critical phase of brain development. Brain Dev (2015).

[8] C. Chen, and S. Tonegawa, Molecular genetic analysis of synaptic plasticity, activity-dependent neural development, learning, and memory in the mammalian brain. Annu Rev Neurosci 20 (1997) 157–84.

[9] S.A. Marik, H. Yamahachi, J.N. McManus, G. Szabo, and C.D. Gilbert, Axonal dynamics of excitatory and inhibitory neurons in somatosensory cortex. PLoS Biol 8 (2010) e1000395.

[10] H. Yamahachi, S.A. Marik, J.N. McManus, W. Denk, and C.D. Gilbert, Rapid axonal sprouting and pruning accompany functional reorganization in primary visual cortex. Neuron 64 (2009) 719–29.

[11] T.C. Thiagarajan, M. Lindskog, A. Malgaroli, and R.W. Tsien, LTP and adaptation to inactivity: overlapping mechanisms and implications for metaplasticity. Neuropharmacology 52 (2007) 156–75.

[12] T.C. Thiagarajan, M. Lindskog, and R.W. Tsien, Adaptation to synaptic inactivity in hippocampal neurons. Neuron 47 (2005) 725–37.

[13] C. Rampon, C.H. Jiang, H. Dong, Y.P. Tang, D.J. Lockhart, P.G. Schultz, J.Z. Tsien, and Y. Hu, Effects of environmental enrichment on gene expression in the brain. Proc Natl Acad Sci U S A 97 (2000) 12880–4.

[14] M. Bayat, M.D. Sharifi, M. Haghani, and M. Shabani, Enriched environment improves synaptic plasticity and cognitive deficiency in chronic cerebral hypoperfused rats. Brain Res Bull 119 (2015) 34–40.

[15] R. Hullinger, K. O’Riordan, and C. Burger, Environmental enrichment improves learning and memory and long-term potentiation in young adult rats through a mechanism requiring mGluR5 signaling and sustained activation of p70s6k. Neurobiol Learn Mem 125 (2015) 126–34.

[16] M.S. Kim, J.H. Yu, C.H. Kim, J.Y. Choi, J.H. Seo, M.Y. Lee, C.H. Yi, T.H. Choi, Y.H. Ryu, J.E. Lee, B.H. Lee, H. Kim, and S.R. Cho, Environmental enrichment enhances synaptic plasticity by internalization of striatal dopamine transporters. J Cereb Blood Flow Metab (2015).

[17] G.I. Irvine, and W.C. Abraham, Enriched environment exposure alters the input-output dynamics of synaptic transmission in area CA1 of freely moving rats. Neurosci Lett 391 (2005) 32–7.

[18] L.R. Stein, K.A. O’Dell, M. Funatsu, C.F. Zorumski, and Y. Izumi, Short-term environmental enrichment enhances synaptic plasticity in hippocampal slices from aged rats. Neuroscience 329 (2016) 294–305.

[19] M. Bose, P. Munoz-Llancao, S. Roychowdhury, J.A. Nichols, V. Jakkamsetti, B. Porter, R. Byrapureddy, H. Salgado, M.P. Kilgard, F. Aboitiz, A. Dagnino-Subiabre, and M. Atzori, Effect of the environment on the dendritic morphology of the rat auditory cortex. Synapse 64 (2010) 97–110.

[20] M. Caleo, D. Tropea, C. Rossi, L. Gianfranceschi, and L. Maffei, Environmental enrichment promotes fiber sprouting after deafferentation of the superior colliculus in the adult rat brain. Exp Neurol 216 (2009) 515–9.

[21] M. Mainardi, A. Di Garbo, M. Caleo, N. Berardi, A. Sale, and L. Maffei, Environmental enrichment strengthens corticocortical interactions and reduces amyloid-beta oligomers in aged mice. Front Aging Neurosci 6 (2014) 1.

[22] M. Mainardi, S. Landi, L. Gianfranceschi, S. Baldini, R. De Pasquale, N. Berardi, L. Maffei, and M. Caleo, Environmental enrichment potentiates thalamocortical transmission and plasticity in the adult rat visual cortex. J Neurosci Res 88 (2010) 3048–59.

[23] J.C. Bennett, P.A. McRae, L.J. Levy, and K.M. Frick, Long-term continuous, but not daily, environmental enrichment reduces spatial memory decline in aged male mice. Neurobiol Learn Mem 85 (2006) 139–52.

[24] S. Koh, H. Chung, H. Xia, A. Mahadevia, and Y. Song, Environmental enrichment reverses the impaired exploratory behavior and altered gene expression induced by early-life seizures. J Child Neurol 20 (2005) 796–802.

[25] D.A. Costa, J.R. Cracchiolo, A.D. Bachstetter, T.F. Hughes, K.R. Bales, S.M. Paul, R.F. Mervis, G.W. Arendash, and H. Potter, Enrichment improves cognition in AD mice by amyloid-related and unrelated mechanisms. Neurobiol Aging 28 (2007) 831–44.

[26] T. Begenisic, M. Spolidoro, C. Braschi, L. Baroncelli, M. Milanese, G. Pietra, M.E. Fabbri, G. Bonanno, G. Cioni, L. Maffei, and A. Sale, Environmental enrichment decreases GABAergic inhibition and improves cognitive abilities, synaptic plasticity, and visual functions in a mouse model of Down syndrome. Front Cell Neurosci 5 (2011) 29.

[27] A.C. Kentner, A. Khoury, E. Lima Queiroz, and M. MacRae, Environmental enrichment rescues the effects of early life inflammation on markers of synaptic transmission and plasticity. Brain Behav Immun 57 (2016) 151–60.

[28] J. Nithianantharajah, and A.J. Hannan, Enriched environments, experience-dependent plasticity and disorders of the nervous system. Nat Rev Neurosci 7 (2006) 697–709.

[29] N.E. Holz, R. Boecker, E. Hohm, K. Zohsel, A.F. Buchmann, D. Blomeyer, C. Jennen-Steinmetz, S. Baumeister, S. Hohmann, I. Wolf, M.M. Plichta, G. Esser, M. Schmidt, A. Meyer-Lindenberg, T. Banaschewski, D. Brandeis, and M. Laucht, The long-term impact of early life poverty on orbitofrontal cortex volume in adulthood: results from a prospective study over 25 years. Neuropsychopharmacology 40 (2015) 996–1004.

[30] K.G. Noble, S.M. Houston, N.H. Brito, H. Bartsch, E. Kan, J.M. Kuperman, N. Akshoomoff, D.G. Amaral, C.S. Bloss, O. Libiger, N.J. Schork, S.S. Murray, B.J. Casey, L. Chang, T.M. Ernst, J.A. Frazier, J.R. Gruen, D.N. Kennedy, P. Van Zijl, S. Mostofsky, W.E. Kaufmann, T. Kenet, A.M. Dale, T.L. Jernigan, and E.R. Sowell, Family income, parental education and brain structure in children and adolescents. Nat Neurosci 18 (2015) 773–8.

[31] P. Tomalski, D.G. Moore, H. Ribeiro, E.L. Axelsson, E. Murphy, A. Karmiloff-Smith, M.H. Johnson, and E. Kushnerenko, Socioeconomic status and functional brain development - associations in early infancy. Dev Sci 16 (2013) 676–87.

[32] B. Boller, S. Mellah, G. Ducharme-Laliberte, and S. Belleville, Relationships between years of education, regional grey matter volumes, and working memory-related brain activity in healthy older adults. Brain Imaging Behav (2016).

[33] S.R. Cox, D.A. Dickie, S.J. Ritchie, S. Karama, A. Pattie, N.A. Royle, J. Corley, B.S. Aribisala, M. Valdes Hernandez, S. Munoz Maniega, J.M. Starr, M.E. Bastin, A.C. Evans, J.M. Wardlaw, and I.J. Deary, Associations between education and brain structure at age 73 years, adjusted for age 11 IQ. Neurology 87 (2016) 1820–1826.

[34] G. Ingram, and A. Will, Global Educational Trends: 1970-2025. FHI 360 Working Paper (2009).

[35] N.A. Badcock, K.A. Preece, B. de Wit, K. Glenn, N. Fieder, J. Thie, and G. McArthur, Validation of the Emotiv EPOC EEG system for research quality auditory event-related potentials in children. PeerJ 3 (2015) e907.

[36] M. Duvinage, T. Castermans, M. Petieau, T. Hoellinger, G. Cheron, and T. Dutoit, Performance of the Emotiv Epoc headset for P300-based applications. Biomed Eng Online 12 (2013) 56.

[37] A.S. Elsawy, S. Eldawlatly, M. Taher, and G.M. Aly, Performance analysis of a Principal Component Analysis ensemble classifier for Emotiv headset P300 spellers. Conf Proc IEEE Eng Med Biol Soc 2014 (2014) 5032–5.

[38] N.A. Badcock, P. Mousikou, Y. Mahajan, P. de Lissa, J. Thie, and G. McArthur, Validation of the Emotiv EPOC((R)) EEG gaming system for measuring research quality auditory ERPs. PeerJ 1 (2013) e38.

[39] P. de Lissa, S. Sorensen, N. Badcock, J. Thie, and G. McArthur, Measuring the face-sensitive N170 with a gaming EEG system: A validation study. J Neurosci Methods 253 (2015) 47–54.

[40] A. Rodriguez, B. Rey, and M. Alcaniz, Validation of a low-cost EEG device for mood induction studies. Stud Health Technol Inform 191 (2013) 43–7.

[41] T. Hwang, M. Kim, M. Hwangbo, and E. Oh, Comparative analysis of cognitive tasks for modeling mental workload with electroencephalogram. Conf Proc IEEE Eng Med Biol Soc 2014 (2014) 2661–5.

[42] T.C. Thiagarajan, M.A. Lebedev, M.A. Nicolelis, and D. Plenz, Coherence potentials: loss-less, all-or-none network events in the cortex. PLoS Biol 8 (2010) e1000278.

[43] C.M. Gray, A.K. Engel, P. Konig, and W. Singer, Synchronization of oscillatory neuronal responses in cat striate cortex: temporal properties. Vis Neurosci 8 (1992) 337–47.

[44] E.S. Kappenman, and S.J. Luck, The effects of electrode impedance on data quality and statistical significance in ERP recordings. Psychophysiology 47 (2010) 888–904.

[45] C.M. Gray, and W. Singer, Stimulus-specific neuronal oscillations in orientation columns of cat visual cortex. Proc Natl Acad Sci U S A 86 (1989) 1698–702.

[46] J.P. Lachaux, N. Axmacher, F. Mormann, E. Halgren, and N.E. Crone, High-frequency neural activity and human cognition: past, present and possible future of intracranial EEG research. Prog Neurobiol 98 (2012) 279–301.

[47] N. Axmacher, F. Mormann, G. Fernandez, C.E. Elger, and J. Fell, Memory formation by neuronal synchronization. Brain Res Rev 52 (2006) 170–82.

[48] G.R. Sutherland, and B. McNaughton, Memory trace reactivation in hippocampal and neocortical neuronal ensembles. Curr Opin Neurobiol 10 (2000) 180–6.

[49] C. Herry, and J.P. Johansen, Encoding of fear learning and memory in distributed neuronal circuits. Nat Neurosci 17 (2014) 1644–54.

[50] E. Thomson, J. Lou, K. Sylvester, A. McDonough, S. Tica, and M.A. Nicolelis, Basal forebrain dynamics during a tactile discrimination task. J Neurophysiol 112 (2014) 1179–91.

[51] K. Dzirasa, R. Fuentes, S. Kumar, J.M. Potes, and M.A. Nicolelis, Chronic in vivo multi-circuit neurophysiological recordings in mice. J Neurosci Methods 195 (2011) 36–46.

[52] D. Gervasoni, S.C. Lin, S. Ribeiro, E.S. Soares, J. Pantoja, and M.A. Nicolelis, Global forebrain dynamics predict rat behavioral states and their transitions. J Neurosci 24 (2004) 11137–47.

[53] D. Parameshwaran, N.E. Crone, and T.C. Thiagarajan, Coherence potentials encode simple human sensorimotor behavior. PLoS One 7 (2012) e30514.

[54] M.S. Fifer, M. Mollazadeh, S. Acharya, N.V. Thakor, and N.E. Crone, Asynchronous decoding of grasp aperture from human ECoG during a reach-to-grasp task. Conf Proc IEEE Eng Med Biol Soc 2011 (2011) 4584–7.

[55] T.H. Grandy, M. Werkle-Bergner, C. Chicherio, M. Lovden, F. Schmiedek, and U. Lindenberger, Individual alpha peak frequency is related to latent factors of general cognitive abilities. Neuroimage 79 (2013) 10–8.

[56] C. Richard Clark, M.D. Veltmeyer, R.J. Hamilton, E. Simms, R. Paul, D. Hermens, and E. Gordon, Spontaneous alpha peak frequency predicts working memory performance across the age span. Int J Psychophysiol 53 (2004) 1–9.

[57] T. Petermann, T.C. Thiagarajan, M.A. Lebedev, M.A. Nicolelis, D.R. Chialvo, and D. Plenz, Spontaneous cortical activity in awake monkeys composed of neuronal avalanches. Proc Natl Acad Sci U S A 106 (2009) 15921–6.

[58] H.A. Al-Nashash, J.S. Paul, W.C. Ziai, D.F. Hanley, and N.V. Thakor, Wavelet entropy for subband segmentation of EEG during injury and recovery. Ann Biomed Eng 31 (2003) 653–8.

[59] V. Jantti, and S. Alahuhta, Spectral entropy‐‐what has it to do with anaesthesia, and the EEG? Br J Anaesth 93 (2004) 150–1; author reply 151-2.

[60] Z. Liang, Y. Wang, X. Sun, D. Li, L.J. Voss, J.W. Sleigh, S. Hagihira, and X. Li, EEG entropy measures in anesthesia. Front Comput Neurosci 9 (2015) 16.

[61] H. Abe, J.N. McManus, N. Ramalingam, W. Li, S.A. Marik, S.M. Borgloh, and C.D. Gilbert, Adult cortical plasticity studied with chronically implanted electrode arrays. J Neurosci 35 (2015) 2778–90.

[62] R.P.K. Sammons, T., Adult plasticity and cortical reorganization afer peripheral lesions. Curr Opin Neurobiol. 35 (2015) 136–141.

[63] H.K. Lee, Whitt J.L., Cross-modal synaptic plasticity in adult primary sensory cortices. Curr Opin Neurobiol. 35 (2015) 119–26.

